# Restoring Protein Glycosylation with GlycoShape

**DOI:** 10.1101/2023.12.11.571101

**Authors:** Callum M Ives, Ojas Singh, Silvia D’Andrea, Carl A Fogarty, Aoife M Harbison, Akash Satheesan, Beatrice Tropea, Elisa Fadda

## Abstract

During the past few years, we have been witnessing a revolution in structural biology. Leveraging on technological and computational advances, scientists can now resolve biomolecular structures at the atomistic level of detail by cryogenic electron microscopy (cryo-EM) and predict 3D structures from sequence alone by machine learning (ML). One technique often supports the other to provide the view of atoms in molecules required to capture the function of molecular machines. An example of the extraordinary impact of these advances on scientific discovery and on public health is given by how structural information supported the rapid development of COVID-19 vaccines based on the SARS-CoV-2 spike (S) glycoprotein. Yet, none of these new technologies can capture the details of the dense coat of glycans covering S, which is responsible for its natural, biologically active structure and function and ultimately for viral evasion. Indeed, glycosylation, the most abundant post-translational modification of proteins, is largely invisible through experimental structural biology and in turn it cannot be reproduced by ML, because of the lack of data to learn from. Molecular simulations through high-performance computing (HPC) can fill this crucial information gap, yet the computational resources, the users’ skills and the long timescales involved limit applications of molecular modelling to single study cases. To broaden access to structural information on glycans, here we introduce GlycoShape (https://glycoshape.org) an open access (OA) glycan structure database and toolbox designed to restore glycoproteins to their native functional form by supplementing the structural information available on proteins in public repositories, such as the RCSB PDB (www.rcsb.org) and AlphaFold Protein Structure Database (https://alphafold.ebi.ac.uk/), with the missing glycans derived from over 1 ms of cumulative sampling from molecular dynamics (MD) simulations. The GlycoShape Glycan Database (GDB) currently counts over 435 unique glycans principally covering the human glycome and with additional structures, fragments, and epitopes from other eukaryotic and prokaryotic organisms. The GDB feeds into Re-Glyco, a bespoke algorithm in GlycoShape designed to rapidly restore the natural glycosylation to protein 3D structures and to predict *N-*glycosylation occupancy, where unknown. Ultimately, integration of GlycoShape with other OA protein structure databases can provide a step-change in scientific discovery, from the structural and functional characterization of the active form of biomolecules, all the way down to pharmacological applications and drug discovery.

## Introduction

The native fold of a protein determines its biological function, regulating mechanisms driving protein-protein and protein-ligand recognition, binding and unbinding events in operational or physiological thermodynamic conditions. The chemical nature of the amino acids and their precise sequence are key determinants towards the correct protein folding and also regulate both structural stability and dynamic properties upon folding. As a result of these crucial roles, protein sequence is strictly safeguarded by genetic encoding to guarantee correct and reproducible biological function. In all forms of life this seemingly rigid, template-driven paradigm is complemented by glycosylation, a remarkably flexible strategy that allows for sequence and structure changes to occur on the fly, reflecting changes in the environmental conditions in different species and states of health and disease, in which proteins are required to operate.

Glycosylation refers to the enzymatically-controlled functionalization of biomolecules with complex carbohydrates, or glycans. It is estimated that about 3-4% of the human genome is devoted solely to encode for mechanisms regulating protein glycosylation^1,2^, underscoring its importance and ubiquity. Glycosylation can occur both, co- and post-translationally through the formation of covalent bonds involving the hydroxyl groups of Ser and Thr sidechains, leading to *O-*glycans, or the amide nitrogen of the Asn sidechain, generating *N-*glycans^3^. Less common, yet highly conserved^4^ modifications include the functionalization at C2 of the Trp indole sidechain with mannose, known as *C-*mannosylation^1,5,6^. The structural complexity of glycans and of glycosylation patterns is unparalleled in Nature, where sequences can range from one monosaccharide to hundreds of units linked through linear and branched arrangements. The glycosidic linkages connecting different monosaccharides add a high degree of flexibility to the structures, which become largely invisible to experiments^7–9^ even in cryogenic conditions. Indeed, glycans are often removed to allow for protein crystallisation^9^. These difficulties in the structural characterization of sugars are further enhanced by the absence of an encoding template, which determines varying degrees of macro and microheterogeneity typical of all glycoconjugates^10–15^.

The sparsity of information on glycan 3D structures, occupation and identity at different sites contributes greatly to our lack of understanding of the many different functions that glycans and glycosylation play in biology. Yet remarkable advances are continuously made in the development of new high-precision tools and techniques^16–26^ that allow us to shed light on different aspects of the glycome. Within this framework, the potentials of glycobioinformatics^27–32^ databases and resources and of high-performance computing (HPC) molecular simulations^7,8^ are extraordinary, especially within a context where information from multiple sources is required to decipher the GlycoCode^33^. Here we introduce GlycoShape (https://glycoshape.org) an open access (OA) web-based platform designed to supplement the missing 3D structural information on glycans and glycoproteins, leveraging on over 1 ms of molecular dynamics (MD) simulations ran in dilute solution at standard conditions of temperature, pressure and salt concentration. A schematic overview of the GlycoShape workflow is shown in **Figure 1**.

**Figure 1.**
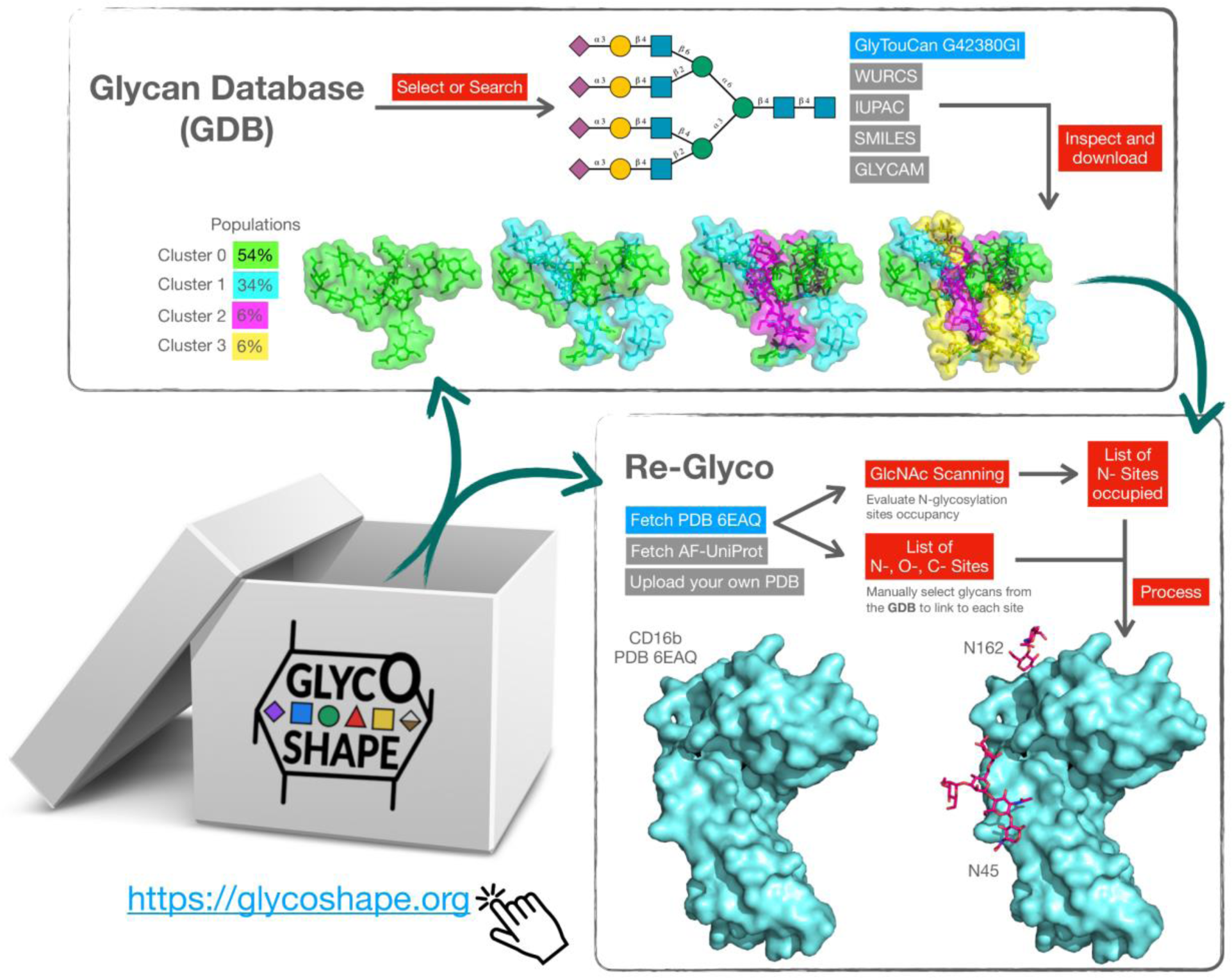
Schematic representation of the GlycoShape workflow (https://glycoshape.org). **Top:** The GlycoShape Glycan Database (GDB) is a repository of glycan 3D structures from 1 ms cumulative sampling though uncorrelated replicas of deterministic molecular dynamics (MD) simulations. Structures can be searched by drawing an SNFG structure with the integrated Sugar Drawer tool^34^ or by searching by text in IUPAC, WURCS, SMILES, GLYCAM and GlyTouCan formats. The GlyTouCan code of the tetra-antennary complex *N-*glycan is shown in the example in blue. The successful search outputs general information on the glycan in the main tab and its 3D structure in the “Structure” tab resulting from clustering analysis of the MD data with the populations (weights) corresponding to each cluster. This information can be downloaded in PDB, CHARMM and GLYCAM formats. **Right:** Re-Glyco allows users to rebuild the 3D structures of glycoproteins to the desired glycoforms by sourcing 3D glycan structures from the GDB and to predict *N-*glycosylation sites occupancy through a bespoke *GlcNAc Scanning* tool. As an example shown here and discussed in the Result section, the CD16b from PDB 6EAQ (in blue) is processed through the GlcNAc Scanning tool, which correctly predicts occupancy of the sequons at N45 and N162^35^. These sites can be *N-*glycosylated with a ‘one-shot’ approach, where the same *N-*glycan structure is chosen to occupy all, as shown in the example, or manually with a site-by-site approach, where the user can select a different *N-*glycan structure at each site.

The GlycoShape glycan database (GDB) currently counts 435 unique glycans to date, predominantly sourced from the human glycome, but with selected examples of glycans structures, fragments and epitopes from other animal, plant, fungi and bacterial species. Each glycan in the GDB is represented through distinct 3D conformers obtained from clustering analysis of the MD trajectories, with both alpha and beta reducing ends and associated weights corresponding to the relative populations of the clusters during sampling. In GlycoShape this 3D information can be used to rebuild glycoproteins to their functional, native state through a bespoke tool, named Re-Glyco (https://glycoshape.org/reglyco), see **Figure 1**. To perform this task the user can fetch protein structures directly from public repositories, namely from the RCSB PDB (https://www.rcsb.org/) with the corresponding PDB ID, and from the AlphaFold Protein Structure Database^36,37^ (https://alphafold.ebi.ac.uk/) with the corresponding UniProt ID, or upload their own PDB file. Re-Glyco currently restores all types of *C-*, *N-* and *O-*glycosylation through a highly efficient genetic algorithm that minimises a loss function accounting for steric clash. Based on the ability of this algorithm to evaluate accessible space and complementarity between the protein landscape and the glycan structure, Re-Glyco can be also used to predict occupancy of *N-*glycosylation sequons with a bespoke tool we named ‘GlcNAc Scanning’. Testing of GlcNAc Scanning against a subset of protein structures (739) from the AlphaFold Database with *N-*glycosylation sites annotated in UniProt as experimentally verified, scores a 92% success rate. All the details on this test are included in Supplementary Material. In the next sections we will present and discuss the construction and features of the GlycoShape GDB and of Re-Glyco, and we illustrate their use and performance also on improving the resolution and prediction of the underlying protein structure through working examples in the Result section. We will then discuss the potentials of these tools for discovery and for advancing our understanding of the many roles of glycans in cell biology and in life science.

### The GlycoShape Glycan Database (GDB)

The conformational complexity of glycans is determined to a large extent by the inherent flexibility of the glycosidic linkages and can be further enhanced by transitions in the structure of the ring, or pucker^38–40^, with probabilities depending on the type of monosaccharide^41^. Furthermore, unlike proteins that extend within a linear arrangement with peptide bonds structurally restrained by electron delocalisation, glycans can be branched and each glycosidic linkage can potentially adopt two different anomeric orientations. The combinatorial explosion of conformers deriving from these considerations would make the structure and dynamics of glycans virtually impossible to characterise. Yet the glycans commonly found in eukaryotic glycoproteins are relatively small, counting 25 or less monosaccharides, with the exception of glycosaminoglycans (GAGs), which for many reasons constitute a class of their own^42^. The reduced molecular size determines that the number of degrees of freedom accessible to the corresponding glycan structures is contained, compared to proteins or nucleic acids, and thus accessible to characterization by exhaustive sampling through MD simulations. More specifically, unlike protein folding the structure and dynamics of isolated saccharides at infinite dilution can be well reproduced from scratch by MD within an all-atoms additive force field^7,8^ approximation, provided sufficient sampling is allowed.

Exhaustive sampling can be achieved by MD through enhanced sampling methods or by deterministic approaches, where the conformational space is explored in a more controlled fashion, through the design of uncorrelated replicas set to cover all the energetically accessible degrees of freedom, which for most saccharides correspond to the glycosidic linkages with the highest flexibility^7,8^. The 3D structure and dynamics of each glycan in the GlycoShape database derives from this approach to deterministic MD sampling, where replicas of 500 ns are designed to sample different and accessible sections of the potential energy surface. For example, a glycan with three (1/2-6) linkages will be analysed through sampling eight distinct conformations, as each linkage can access only two gauche (*gt* and *gg*) conformations of the three theoretically possible^7,8^. These eight uncorrelated MD replicas will generate a total of 4 μs of cumulative sampling data. The chemical nature of the glycosidic linkages determines that high energy barriers, difficult to overcome at room temperature, are uncommon and in most cases different conformers are seen to rapidly interconvert though this MD approach. These conformers are in equilibrium at room temperature and their ensemble defines the glycan structure. Therefore, the dominant conformation of a glycan can change depending on how the environment stabilises each conformer differently, shifting the equilibrium^43^. The GlycoShape platform provides all the representative structures of the conformers at equilibrium in solution with corresponding weights derived from the MD trajectories through our Glycan Analysis Pipeline (GAP), see **Figure 2**.

**Figure 2:**
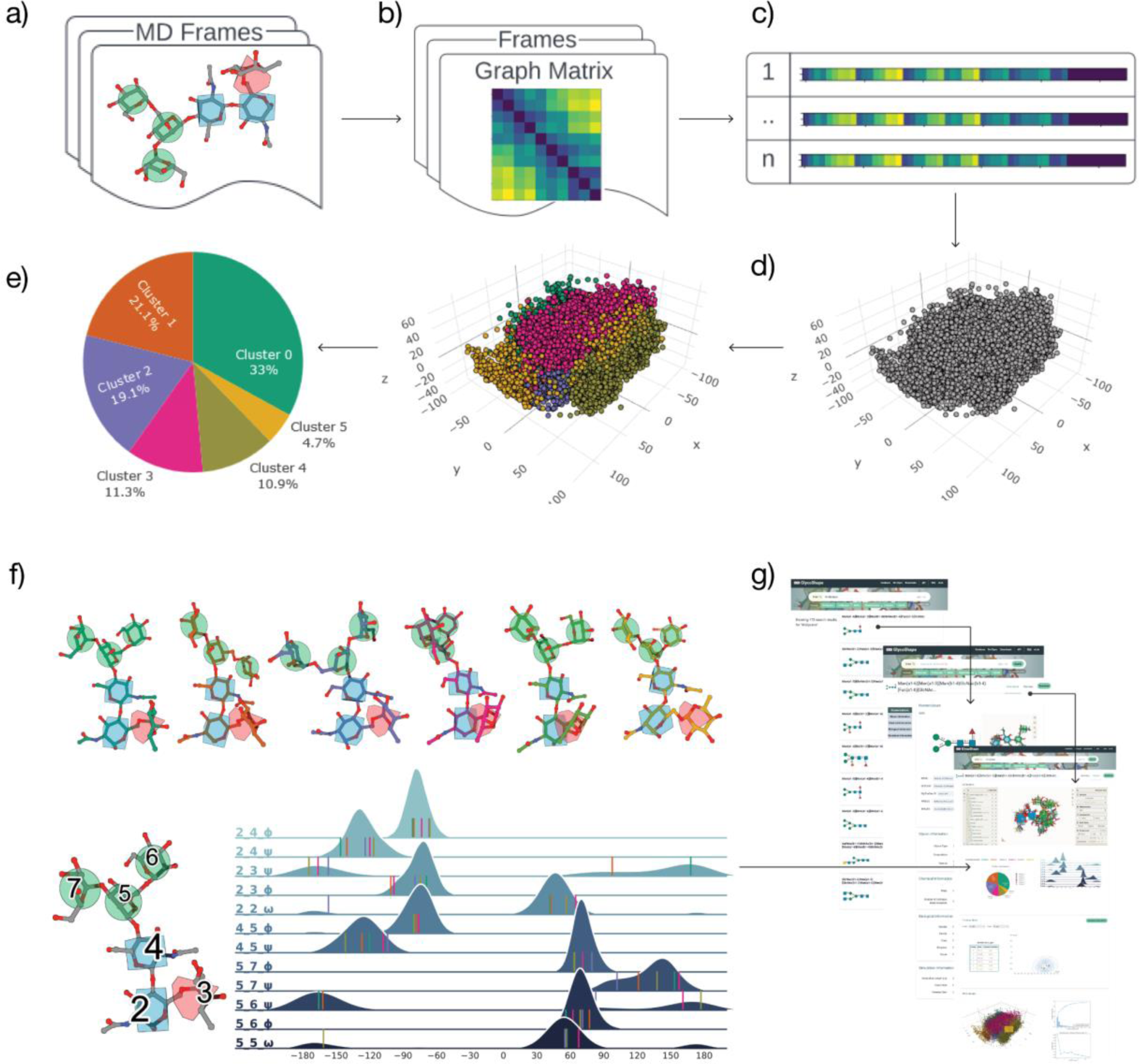
Schematic overview of Glycan Analysis Pipeline (GAP) used to build the GlycoShape Glycan Database (GDB). **Panel a)** Multiple uncorrelated replica molecular dynamics (MD) simulations are performed for each glycan in the GDB, to comprehensively sample its structural dynamics. The resulting MD frames are then transformed into a graph matrix representation, as depicted in **Panel b)**, simplified by flattening the lower half as shown in **Panel c).** This step enables a dimensionality reduction via principal component analysis (PCA), shown in **Panel d)**. These data are clustered by Gaussian Mixture Model (GMM) and the results of which are displayed in terms of cluster distributions, see **Panel e)**. **Panel f)** Representative 3D structures for each cluster are selected based on KDE maxima, along with comprehensive torsion angle profiles for the highest populated clusters, showing the wide breadth of the conformational space covered by GAP. **Panel g)** Structures derived from GAP are clearly presented on the GlycoShape GDB web platform, in addition to biological and chemical information.

In the GAP for each glycan, individual MD trajectories from the uncorrelated replicas are integrated into one dataset. Each frame of this cumulative trajectory is transformed into a graph matrix, from which we derive a one-dimensional array by flattening the matrix’s lower half, see **Figure 2b** and **2c**. This transformation captures all geometric data, setting the stage for a conformational landscape representation. To extract representative glycan structures, these data points are organised into distinct clusters. Given the high dimensionality of the system, we run a dimensionality reduction via principal component analysis (PCA), which leads to a three-dimensional representation of the conformational landscape, see **Figure 2d**. These data are clustered using a Gaussian Mixture Model (GMM), where the optimal number of clusters is determined by the silhouette score, typically between two and ten clusters, often reaching an optimum at five, see **Figure 2e**. The selection of representative 3D structures for each cluster is not merely the centroid, which would correspond to a mean position lacking in conformational specificity. Instead, the most statistically significant conformation for each cluster is better defined through a kernel density estimation (KDE) analysis, which ensures a more accurate representation of the conformational space within each cluster, see **Figure 2f**. Based on GAP users are able to access a wealth of structural information for each glycan structure, which includes not only representative conformational structures, but also their population frequency during extensive MD sampling and detailed information on the ϕ, ψ, and ω torsion angles values. Additionally, the GlycoShape GDB complements this broad array of structural data with biological and chemical data of broad interest, in a clear and understandable format, see **Figure 2g**. More specifically, additional information includes alternative naming formats for each glycan, together with chemical and biological information directly sourced from different glycoinformatics repositories^27,44,45^, as detailed in **Table S.1**. Further details on the design and implementation of the GDB are available in Supplementary Material.

### Rebuilding Glycoproteins with Re-Glyco

Understanding the functional state of a protein requires restoring its co- and post-translational modifications (PTMs), which can affect its structural stability, dynamics and recognition/interaction ability. Re-Glyco is an algorithm implemented in GlycoShape, designed to restore glycosylation to protein 3D structures. The occupancy and specific type of glycosylation results from a complex equilibrium involving hundreds of glycosylhydrolases (GHs) and glycosyltransferases (GTs) operating through the secretory pathway^1,2,46^, within a sequence with priorities still poorly understood. As the protein folds, GHs and GTs enzymatic functions are regulated not only by the levels of expression of the key enzymes, but also by the physical accessibility of the glycosylation site^10,14,15,47,48^. Based on these considerations, restoring glycosylation onto a folded protein to produce representative 3D structures of a glycoprotein requires, 1) access to exhaustive information on the type of glycan at each site^11,13,49^, together with the conformational space accessible to those glycans^7,43^, and 2) an algorithm able to assess the structural complementarity between these glycan 3D structures and the protein rugged landscape surrounding the glycosylation site^43,47,48^.

Re-Glyco sources glycans 3D structures from the GlycoShape GDB. The conformational ensemble for each glycan is represented by the structures corresponding to the KDE-max of each cluster, obtained from the analysis of the MD simulations. Within the assumption of an exhaustive sampling, these 3D structures represent the conformational space accessible to the glycan at 300 K and 1 atm. As discussed in earlier work^43^, the protein structure surrounding the glycosylation site modulates the conformational equilibrium corresponding to the glycan unlinked in solution, to favour the structure with optimal complementarity to the protein 3D landscape. For sites that are spatially accessible, the conformational equilibrium of the linked glycan is identical to the equilibrium corresponding to the unlinked glycan in solution^43^.

The complementarity between the protein landscape surrounding glycosylation sites and the representative structures from each cluster is evaluated by Re-Glyco through a genetic algorithm designed to minimise a loss function F(P, G, φ, ψ), see **Figure 3a**, designed to evaluates steric hindrance gauged in terms of linear distance between all the protein (P) atoms and all the glycan (G) atoms (except for the terminal atom at the reducing end) with a minimum threshold value set to 1.7 Å, corresponding to the van der Waals radius of C. In the loss function F(P, G, φ, ψ), φ and ψ are torsion angles values defining the conformation of the bond to the protein, see **Figure 3a**. The values of φ, ψ have been sourced from Privateer^52^ where available, or obtained from MD simulations used to complete the data set. The Re-Glyco workflow starts by testing the highest populated glycan structure, namely cluster 0 (G0), searching through the predetermined values of ϕ and ψ torsions. Re-Glyco minimises the loss function F(P,G,ϕ,ψ) over a population of 128 torsion sets (φ, ψ) across 8 generations. If clashes persist, the search for P-G(0) is aborted and Re-Glyco shifts to testing the second highest populated conformer from the GlycoShape GDB, i.e. cluster 1 (G1), and so on. If none of the available glycan conformations fits the protein site, Re-Glyco progresses by trying to solve the P-G(0-4/5) interaction that corresponds to the lowest steric hindrance through a ‘wiggle’ process. During the wiggle phase, torsion angle values are adjusted through random moves within a range of 10°, corresponding to the lowest range of the standard deviations commonly associated to the dynamics of glycosidic linkages at 300 K^7,8^. This process is repeated up to 40 times. Further details on the algorithm design and implementation are available in Supplementary Material.

**Figure 3.**
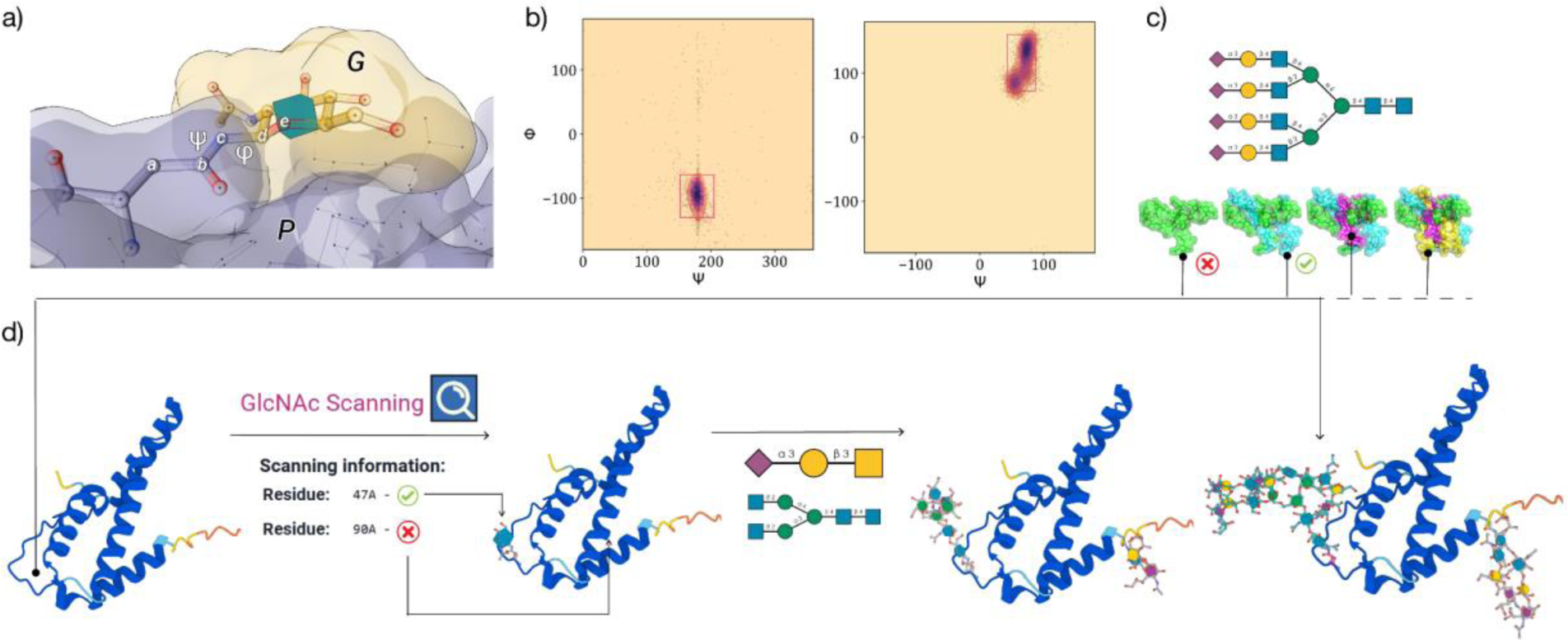
**Panel a)** Definition of the φ and ψ torsion angles, with corresponding atoms labelled a to e, determining the conformation of the linkage between the protein (P shown in grey) sidechain and the reducing end of the glycan (G shown in yellow). **Panel b)** Heat maps showing preferential conformation of the φ, ψ torsions between Asn-b-GlcNAc and Thr-a-GalNAc. **Panel c)** 2D SNFG structure and (below) 3D structures of the tetra-antennary fully a3-sialylated *N-*glycan from the clustering analysis shown in **Figure 1. Panel d)** Schematic representation of the Re-Glyco workflow applied to the reconstruction of human Interleukin-5 (IL5; Uniprot P05113). In agreement with the annotation^50^, GlcNAc scanning identifies only the N47 sequon as potentially occupied. Accordingly N47 can be functionalized with more elaborate structures through a ‘one-shot’ glycosylation, where also T22 can be functionalized with a sialylated core1 O-glycan. Highly complex glycosylation at N47 and alternative O-glycosylation structures can be selected by sourcing directly from the GDB through the Advanced (Site-by-Site) Glycosylation tool, as shown on the alternative IL5 glycoform on the right-hand side. Molecular rendering with Mol* Viewer^51^ and statistical analysis and heat maps created with matplotlib (https://matplotlib.org/).

The ability of Re-Glyco to resolve steric clashes can be used within GlycoShape also to assess the potential occupancy of *N-*glycosylation sites through an implementation we called ‘GlcNAc Scanning’, see **Figure 3d**, to paraphrase the widely used Ala Scanning approach. Where the *N-* glycosylation state of a protein is not known, the user can choose to perform an initial GlcNAc Scanning, where Re-Glyco will try to fit a single GlcNAc monosaccharide into all the NXS/T sequons in the protein of choice with the protocol described above. The process outputs a list of sequons that passed the test, marked with a simple ‘yes’ or ‘no’ label. The sequons labelled with ‘yes’ can be further *N-*glycosylated with longer glycans, by one-shot *N-*glycosylation, where the same *N-*glycan is chosen for all sites. Where different types of glycans are required or desired, the user can manually choose the desired glycosylation at each site predicted to be occupied with the Advanced (Site-by-Site) Glycosylation module. Alternatively, where information on the occupancy of the protein glycosylation sites is available through UniProt, the user can follow directly the given annotation and glycosylate the sites highlighted as occupied in one shot. Please note that this information in UniProt may not always correspond to the functional glycosylation state of the protein^53^, unless supported by published references.

## Results

In this section we present the results of two test cases where GlycoShape was used to restore the biologically active structure of glycoproteins. For this purpose we chose to present particularly challenging scenarios, where the GDB and Re-Glyco are instrumental not only to rebuild glycosylation, but also to flag flaws in the protein structure, or shortcomings of its use as a model. Within this framework, we describe how the occurrence of ‘false positives’ and ‘false negatives’ can lead the user to refine the structural data, either own or sourced from a repository. As a general metric, the Re-Glyco GlcNAc Scanning tool predicts correctly 92% of the *N-*glycosylation sites, based on 739 proteins for which experimental structures are available and with *N-*glycosylation sequons annotated in UniProt as verified by experiments.

### Re-glycosylation of CD16b

As a challenging working example, here we present the performance of Re-Glyco in detecting and rebuilding the functional form of the Fc γ receptor III (CD16), the primary receptor responsible for antibody-dependent cell-mediated cytotoxicity (ADCC)^35,54,55^, see **Figure 4**. CD16 exists in two forms, CD16a and CD16b, that share 97% sequence identity. A crucial mutation from Gly to Asp at position 129 (G129D) in CD16b was found to dramatically reduce binding affinity to the IgG-Fc relative to CD16a^35^. CD16a and CD16b share five of the six *N-*glycosylation sequons^56^, and only one *O-*glycosylation site was detected in recombinant CD16a^57^. In agreement with this profile and for the purpose of this example we chose not to introduce *O-*glycans, but the user can do that by selecting the site manually in the Advanced (Site-by-Site) Glycosylation tool and by adding the desired *O-*glycan structure from the GDB.

**Figure 4.**
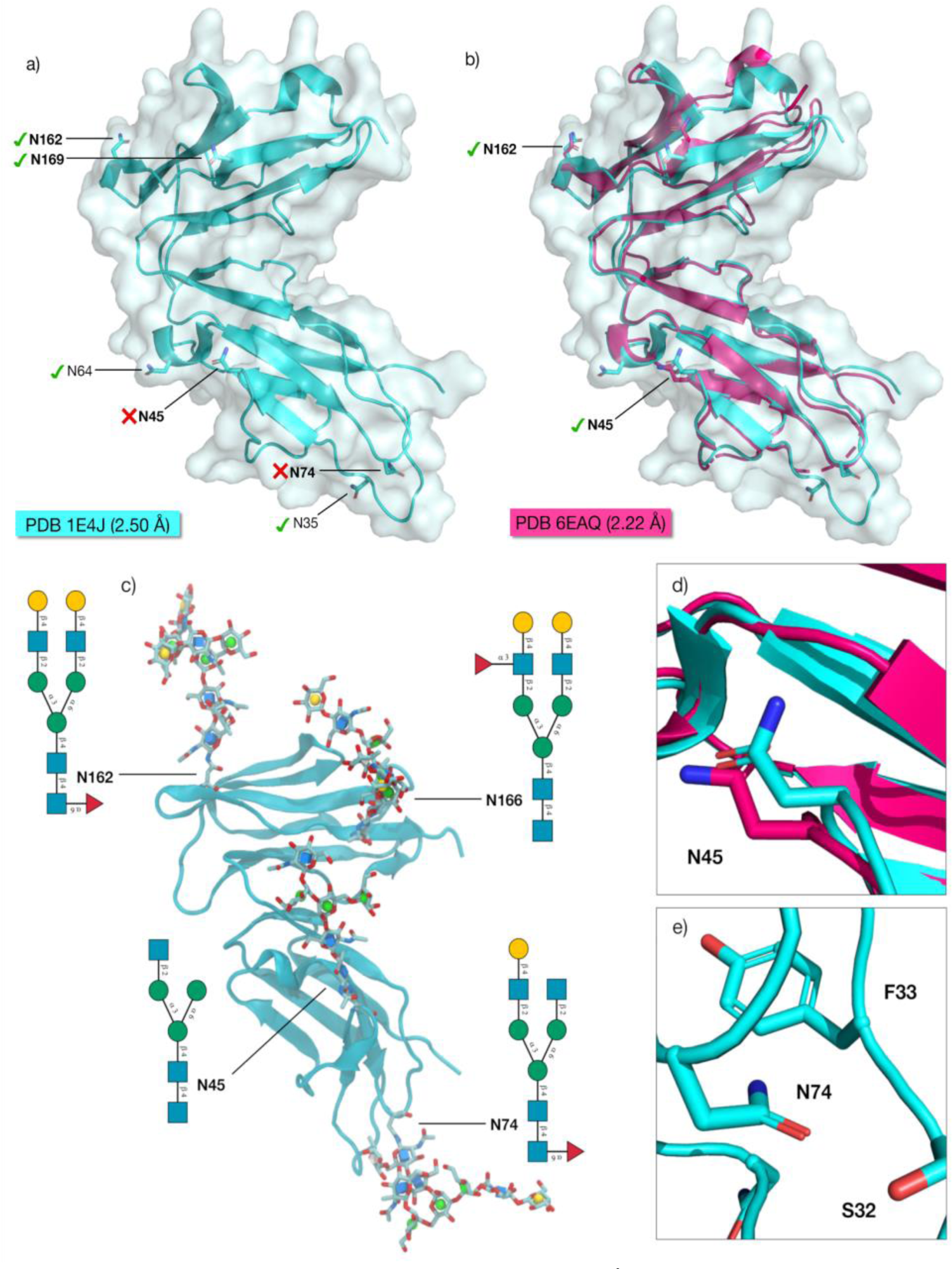
**Panel a)** Structure of the CD16b (PDB 1E4J, resolution 2.50 Å) with all the *N-*glycosylation sequons labelled. The bold labels indicate that the sequons are occupied in neutrophil-bound CD16b^56^. The green check marks indicate the sequons predicted to be occupied by the Re-Glyco GlcNAc-Scanning tool, while the red x-marks indicate the sequons that Re-Glyco deems unoccupied. **Panel b)** Structural alignment of the CD16b (PDB 1E4J) in cyan with the CD16b (PDB 6EAQ, resolution 2.22 Å) in magenta. Both sequons occupied in PDB 6EAQ, namely N45 and N161, are correctly predicted as occupied by Re-Glyco. **Panel c)** Structure of the CD16b (PDB 1E4J) modified by swapping OD1 and ND2 coordinates and alternative rotameric orientation of the N74 sidechain N-glycosylated by Re-Glyco with a different selection of *N-*glycan structures from the GlycoShape GDB at each site. **Panel d)** Close-up view of the OD1 and ND2 orientation of N45 in the CD16b from PDB 1E4J (cyan) and from PDB 6EAQ (magenta). **Panel d)** Close-up of the orientation of the N74 sidechain in the CD16b from PDB 1E4J. The distance between the CG of N45 and the CA of F33 is 4.7 Å. Rendering of the 3D structures in panels a,b, d and e and rotamer search performed with pymol (https://pymol.org/2/). Rendering of 3D structure in panel c with VMD (https://www.ks.uiuc.edu/Research/vmd/) and N-glycans 2D structures with DrawGlycan-SNFG (http://www.virtualglycome.org/DrawGlycan/).

The crystal structure of the soluble CD16b in solution (PDB 1E4J) at 2.5 Å resolution, bears six *N-*glycosylation sequons (allotype NA2), namely N38, N45, N64, N74, N162 and N169, see **Figure 4a**. To note, Re-Glyco sources the residue numbers from the uploaded structure, so in the case of PDB 1E4J residues are indicated with a −3 shift, namely N35, N42, N61, N71, N159 and N166. Here we will use the former standard numbering above to avoid confusion. In the case of neutrophil-bound CD16b an accurate *N-*glycosylation profile is available, not only indicating site occupancy, but also the type of *N-*glycosylation found at each site^56^. According to these data^56^, only four out of the six *N-*glycosylation sites are occupied in the neutrophil-bound human CD16b, namely N45, N74, N162 and N169. The results of the GlcNAc Scanning of the CD16b structure from PDB 1E4J indicates that a single Asn-b-GlcNAc can fit at positions N38, N64, N162 and N169, missing two of the occupied *N-*glycosylation sites at N45 and N74 (two false negatives), and indicating as potentially occupied the unoccupied N38 and N64 (two false positives). As described below, a more in depth analysis reveals the source of these apparent discrepancies, while informing the user about potential flaws in the protein structure.

False negatives in Re-Glyco can flag the wrong orientation of the Asn sidechain, which in the case of N45 depends on the wrong assignment of the positions of ND2 vs. OD1, see **Figure 4a**. The correct orientation of the ND2 and OD1 at N45 is confirmed by the structure of the CD16b in PDB 6EAQ, see **Figure 4b** and **4d**, which was solved with *N-*glycans at N45 and N162, while sequons at N38, N64, N74 and N169 were mutated to QXS/T^35^. As the correct assignment of coordinates to ND2 and OD1 is largely arbitrary, in the case of a clash Re-Glyco includes a tool that allows the user to swap the coordinates of ND2 with OD1 (https://glycoshape.org/swap). The modified PDB can then be screened again. In more complex cases, false negatives can indicate a wrong orientation of the whole Asn sidechain, as shown by the case of N74, see **Figure 4e**. Indeed in PDB 1E4J the N74 sidechain is facing a loop section between S32 and F33, which clearly hinders the addition of a GlcNAc. No further support of the actual orientation of the N74 sidechain is available, as the whole region is not resolved in PDB 6EAQ, see **Figure 4b**, suggesting a high degree of conformational disorder. In this case the clash can be solved by choosing an alternative orientation of the Asn sidechain and reloading the modified PDB structure. Re-Glyco does not provide an automatic rotamer search, as we think that modifying the protein structure, even if to a minor degree, should not be done lightly, i.e. unless informed by expert knowledge or advice.

In regard to false positives, Re-Glyco can predict occupancy of unoccupied sequons when the protein structure under analysis is only part of a more complex system, for example a single domain of a more complex protein, or an isolated monomer of a dimer/trimer/multimer, or, as in the case of CD16b, a transmembrane receptor severed from the membrane. When only part of the protein is processed through GlcNAc Scanning, *N-*glycosylation sites that are buried in the functional form can be accessible. Accordingly, the predicted occupation of the highly accessible N38, see **Figure 4**, can be denied by its proximity to the membrane in the neutrophil-bound form of CD16b^56^. Indeed, *N-*glycosylation at N38 was detected in the recombinant receptor^56^, which corresponds more closely to the isolated structure model processed by Re-Glyco. Analogously, the false positive at N64 where occupancy is suggested by Re-Glyco, but not experimentally verified in neutrophil-bound CD16b^56^, is not straightforward to rationalise. As shown in **Figure 4**, N64 is highly accessible and indeed it is shown to be occupied with highly processed complex *N-*glycans in the soluble form of CD16b^56,58^, suggesting again that the complexity of the environment of which the protein is part, can significantly affect occupancy. To note N64 is the only *N-*glycosylation sequon not conserved between CD16a and CD16b, and it is only present in the CD16b allotype NA2^56^. In summary, within the limitations of an isolated PDB structure representing a membrane bound CD16b receptor and based on the original assignment of the Asn sidechain orientation in the deposited PDB, Re-Glyco provides an accurate prediction of the *N-*glycosylation profile of CD16b allotype NA2. The occupied sequons, can be functionalized with *N-*glycans to match the neutrophil-bound CD16b glycoproteomics profile^56^, as shown in **Figure 4c**.

### Re-Glyco on AlphaFold Structures

In earlier work^59^ some of us and others discussed the potentials of protein 3D structures derived from ML to be completed with the necessary PTMs required for function. More specifically, we argued that protein structures from the AlphaFold Protein Structure Database (https://alphafold.ebi.ac.uk/) can be readily completed with non-protein elements^59^, as predicted features are learned from biologically active protein structures, which may have cofactors, metals, ligands and often PTMs. In the case of glycosylation, while the structural information on the glycan is often incomplete and seldom erroneous^9^ due to the limitations discussed earlier, we find that the structure of the aglycone obtained from ML is suited for functionalization^59^.

In this work we investigated this point further by testing 3,415 proteins from the AlphaFold Protein Structure Database with glycosylation sites annotated in Uniprot. For the reconstruction of a total of 12,789 *N-*glycosylation sites with the basic complex heptasaccharide A2 from the GDB (GlcNAc(b1-2)Man(a1-3)[GlcNAc(b1-2)Man(a1-6)]Man(b1-4)GlcNAc(b1-4)GlcNAc; GlyTouCan ID G88876JQ), Re-Glyco reports no clashes in 10,821 cases (85%) and unresolvable clashes in 1,968 cases (15%). We find that when the clash occurs between the glycan and spatially neighbouring loops with a predicted conformation ranked within the lowest and highest range of the predicted per-residue local distance difference test (pLDDT) scores^36,60,61^, a clash resolution is generally achieved by the use of Re-Glyco alone, see **Figure 5a**. In case of clashes with regions predicted with a medium range confidence score, namely 90 > pLDDT >65 see **Figure 5a**, the resolution may require a direct (manual or otherwise) intervention, which often simply involves the selection of an alternative orientation of the sidechain of the aglycone or of the clashing residue, as discussed in the previous subsection. A pLDDT value below 70 is reported to indicate a lower degree of confidence in the AlphaFold 3D structure prediction^61^, yet the results of a community-based exhaustive evaluation^62^ suggest that the user should feel justified in selecting alternative sidechain orientations for residues with an associated pLDDT score < 90.

**Figure 5.**
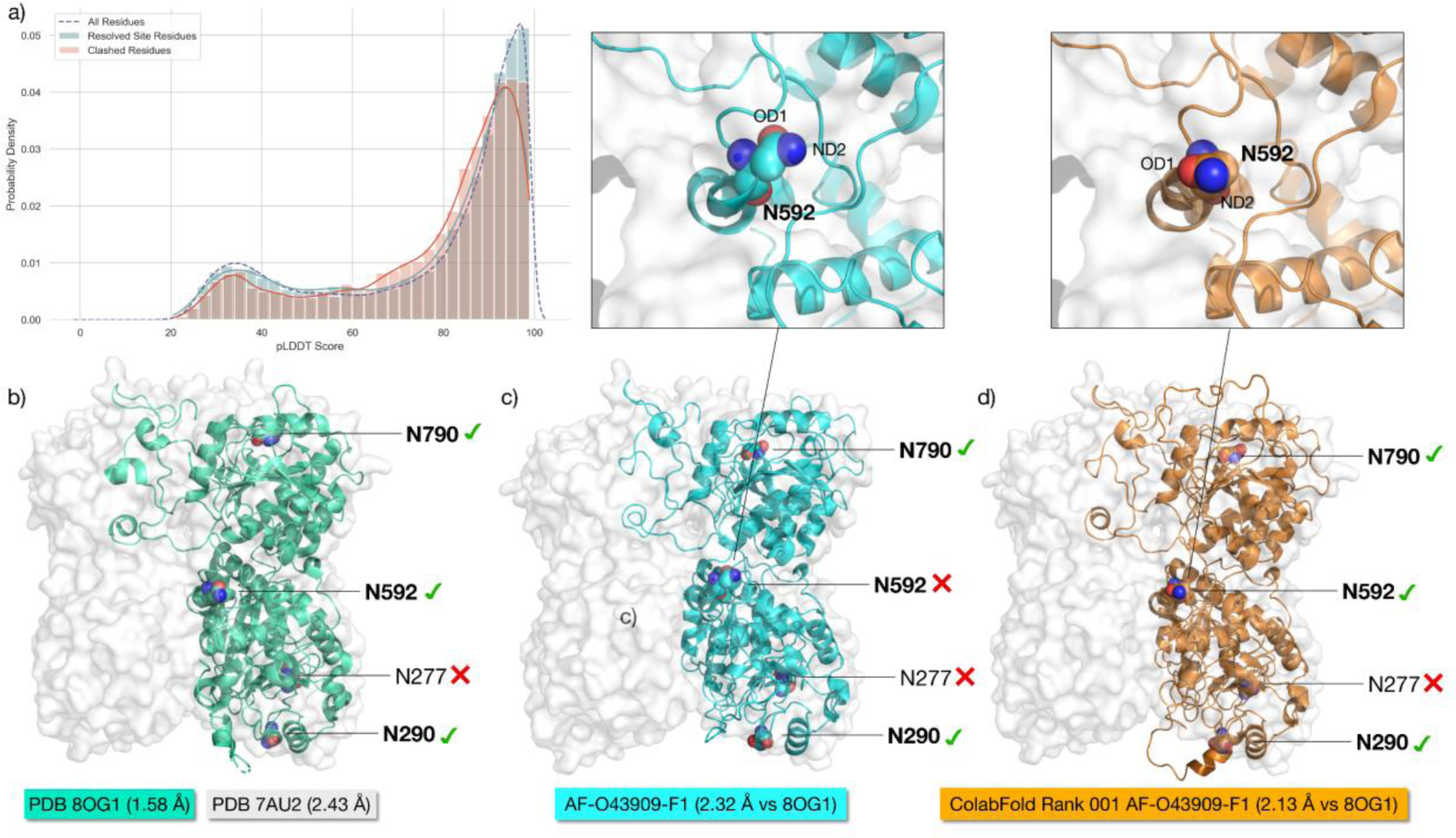
**Panel a)** Histogram analysis of the distribution of pLLDT scores of residues clashing during the GlcNAc Scanning of 3,415 proteins from the AlphaFold Protein Structure Database, with a total of 12,789 glycosylation sites annotated in Uniprot. The distribution of pLDDT values for all residues in the protein tested is shown with a dotted line. The distribution of the pLDDT values for the residues clashing with the GlcNAc during GlcNAc Scanning, where clashes were resolved by Re-Glyco is shown with powder blue histograms and a blue line. The distribution of the pLDDT values for the residues where clashing was not resolved by Re-Glyco and for the residues in the immediate vicinity (± 2) is shown with rose histograms and a red line. **Panel b)** 3D structure of the EXTL3 monomer from PDB 8OG1 (green cartoons) represented within the homodimer from PDB 7AU2 (white surface). The resolution for each structure is shown in the labels. Asn residues within sequons are shown with van der Waals (vdW) spheres, where N atoms are in blue and O atoms are in red. The *N-*glycosylation sequons known to be occupied are shown in bold. A green checkmark indicates that the site is predicted to be occupied by Re-Glyco, while a red cross mark indicates that the site is predicted to be unoccupied by Re-Glyco due to major steric clashing. **Panel c)** 3D structure of the EXTL3 monomer from AlphaFold (AF-O43909-F1) shown in cyan. The lowest confidence loops and termini are removed from the image for clarity. The all-atom RMSD vs PDB 8OG1 is shown in the label. A close-up view of the sidechain orientation of Asn 592 is shown in the panel above, where the clash with a spatially neighbouring loop prevents functionalization. **Panel d)** 3D structure of the EXTL3 monomer from ColabFold (Rank 001) shown in orange. A close-up view of the sidechain orientation of Asn 592 is shown in the panel above, showing the alternative orientation that allows for functionalization. Molecular representation with pymol (https://pymol.org/2) and statistical analysis and rendering created with matplotlib (https://matplotlib.org/).

A conformational search aimed at finding the optimal orientation of a residue sidechain may not be a strategy amenable to all users. To this end we show here an example of how clashes encountered during the reconstruction of glycoproteins from the AlphaFold Database can be often resolved by using ColabFold^63^, as shown in **Figure 5**. To illustrate this case, we selected the structure of the human exostosin-like 3 (EXTL3), one of the five exostosins^64^ known to contribute to the synthesis of the heparan sulphate backbone in the Golgi^65^. EXTL3 not only has an AlphaFold structure (AF-O43909-F1), but was also resolved by cryo-EM^66^ (PDB IDs 7AU2 and 7AUA) and by X-ray crystallography^67^ (PDB IDs 8OG1 and 8OG4). In **Figure 5** we show the structures of the EXTL3 monomers from PDB 8OG1 (panel 5b), from AlphaFold (panel 5c) and from ColabFold (panel 5d) within the silhouette of the homodimer based on the solvent accessible surface from PDB 7AU2. *N-*glycosylation of human EXTL3 was experimentally confirmed^68^ at sites N290, N592 and N790, while the sequon at N277 is unoccupied. GlcNAc Scanning with Re-Glyco of the experimental structures leads to the correct identification of all occupied sequons, and also to the correct rejection of the sequon at N277. GlcNAc Scanning of the AlphaFold structure reports a false negative at N592 due to the wrong orientation of the Asn 592 sidechain, see **Figure 5c**. This clash is resolved by using one of the five structures provided by ColabFold^63^, namely Rank 001, which in this case gives an orientation of the N592 sidechain suitable for functionalization, see **Figure 5d**.

## Discussion and Conclusions

The reconstruction of the CD16b and of the EXTL3 structures described in the previous section highlights some of the features, potentials and unique capabilities of GlycoShape that we believe will help glycobiologists, structural biologists and the broader scientific community to easily restore glycoproteins to their functional form. MD simulations can now produce highly reliable results that can be directly compared to experiments, enabling researchers to understand biological function at the molecular level of detail.

*N-*glycosylation is among the most common types of protein PTMs, instrumental to protein folding and structural stability^46,69–71^, as well as to mediating countless interactions between the cell and its environment in health and disease. *N-*glycans can also function as the first port of entry for pathogen infection^72^ and are determinant to viral fitness and evasion^47,73–77^.

Understanding the many roles of *N-*glycosylation requires the assessment of occupancy at each sequon and of the type of glycosylation at each occupied site. This information can be obtained by glycoproteomics, which to date remains the gold-standard for protein glycoanalysis. Yet, glycoproteomics profiles are not available for all glycoproteins and require dedicated studies. The GlcNAc Scanning tool in Re-Glyco provides an additional, rapid screen of the accessible volume at each *N-*glycosylation sequon and reports the potential occupation of the site by a single β-linked GlcNAc with a simple yes and no answer. This is in line with the determinant role of site accessibility as a determinant for *N-*glycosylation^10,11,14,47,78^, along with other considerations, further supported by the correct identification of 92% of experimentally confirmed *N-*glycosylation sites in a subset of 739 proteins from the AlphaFold database. Re-Glyco provides the means to functionalize the sites predicted to be occupied with more elaborate glycan structures to match the desired profile. This selection should be always guided by glycoproteomics data, where available. Nevertheless, we are currently working on strategies to provide users with an informed choice of potential glycosylation profiles corresponding to protein expression in different cell lines.

In the case of *O-*glycans, prediction of site occupancy is still one of the most difficult challenges in glyco-bioinformatics. Functionalization with *O-*GalNAc glycans, which is the most abundant type of *O-*glycosylation of proteins^79^ and that is inherently linked to their function^80,81^, does not follow a precise amino acid sequence and is frequently encountered in highly flexible protein regions. Therefore, an approach based on sampling 3D space accessibility at potential *O-* glycosylation sites, would not help. At this stage users interested in reconstructing *O-* glycosylated sites with GlycoShape are advised to follow profiles obtained from glycoproteomics data and to rebuild the desired glycoform through a *site-by-site* approach in Re-Glyco. Nevertheless, a *one-shot* glycosylation is available also for *O-*GalNAc glycosylation, where the *O-*glycan selected as default is a sialylated core1 type.

In the result section we have shown different cases where GlycoShape is not only useful to restore glycoprotein structures to their natural and functional form, but also how it provides an efficient strategy to identify minor flaws in the protein structure. Indeed, GlcNAc scanning can indicate the position of the Asn OD1 and ND2 atoms that experimentally is rarely attainable and can also inform the choice of the Asn sidechain orientation. Where the protein structure requires a direct alteration to allow glycosylation with Re-Glyco to proceed, GlycoShape does not offer a rotamer search tool as we believe that protein structure modifications are not always justified and should be carefully guided by expert advice. Yet, one can argue that for ML predicted structures rotamer optimisation should always be allowed. Our analysis of a subset of glycosylated proteins from the AlphaFold database, shows that Re-Glyco finds unresolvable clashes more often with residues with a mid-range pLDDT score. We also show that these clashes can be resolved by using alternative outputs, which are easily obtained through ColabFold^63^ in the absence of dedicated computing resources. Ultimately, the results in this work confirm an argument some of us presented earlier^59^ as well as recent work from the developers of RoseTTAFold^82,83^, that protein 3D structures from ML can be readily glycosylated and also used to predict glycosylation site occupancy, where this depends on 3D structure accessibility. For these reasons we believe that a direct integration of GlycoShape within existing structural databases or ML-workflows would greatly help users to include glycosylation in their work and promote a better understanding of the many roles of glycans in the biology of health, disease and in life science.

## Data availability

The Glycan Database (GDB) is publicly accessible at https://glycoshape.org. In addition, the molecular dynamics (MD) simulations, which were used to curate the GDB, are available for download at https://glycoshape.org/downloads.

## Code availability

The source code for the clustering methodology can be accessed at the following GitHub repository: https://github.com/Ojas-Singh/GlycanAnalysisPipeline. For the glycoprotein builder, the code is available at https://github.com/Ojas-Singh/Re-Glyco.

## Authors Contributions

CMI developed and curated the GlycoShape GDB. OS developed the clustering analysis methodology, Re-Glyco and GlycoShape website. CMI, OS, SDA, CAF, AMH, AS, BT ran all MD simulations used to build the GlycoShape GDB. EF designed and developed the GlycoShape project and overviewed its development. CMI, OS and EF analysed the data and wrote the manuscript.

## Acknowledgements

All authors would like to thank all colleagues and friends who carefully tested the beta version of GlycoShape, providing us with feedback, bug reports, and crucial information that led us to finalise the current release. The Science Foundation of Ireland (SFI) Frontiers for the Future Programme is gratefully acknowledged for financial support of CMI postdoctoral training and of SDA and OS postgraduate training (20/FFP-P/8809). The Science Foundation of Ireland (SFI) Centre for Research Training in Data Science (www.data-science.ie) is gratefully acknowledged for financial support of AS and BT postgraduate training (18/CRT/6049). The opinions, findings, and conclusions or recommendations expressed in this material are those of the author(s) and do not necessarily reflect the views of the Science Foundation of Ireland. CAF acknowledges the Irish Research Council (IRC) for funding through the Government of Ireland Postgraduate Scholarship Programme (GOIPG/201912212). We gratefully acknowledge ORACLE for Research for the generous allocation of computational resources and Richard Pitts and Mike Riley at ORACLE for Research for their feedback, encouragement, and support. CMI, CAF, AMH and EF acknowledge the Irish Centre for High-End Computing (ICHEC) for generous allocation of computational resources. EF gratefully acknowledges the contributions to the GDB derived from the work of Irene Cuxart Sanchez from Prof Carme Rovira Virgili’s research group, during her placement in EF lab at Maynooth University. AMH is currently a postdoctoral fellow at University College Dublin (UCD).

