## Supplementary Material for "Restoring Protein Glycosylation with GlycoShape"

#### Contents

**1. GlycoShape Website** ... **3**
1.1. Cross referenced data ... 3
1.2. Frontend and APIs ... 4

**2. Free Glycan Simulation Protocol** ... **5**

**3. Glycan Analysis Pipeline** ... **7**
3.1. Molecule representation ... 7
3.2. Dimentionality reduction ... 7
3.3. Clustering algorithm and number of clusters ... 8
3.4. Representative structure from clusters KDE maxima ... 8

**4. Re-Glyco: a glycoprotein builder** ... **10**
4.1. Glycoprotein torsion information ... 10
4.2. $P$, $G$ and fitness function $F_{\{P,G,\varphi,\psi\}}$ ... 13
4.3. Wiggle algorithm for alternate conformation within a cluster ... 14
4.4. Genetic optimization algorithm for finding optimal $\varphi$ and $\psi$ ... 16
4.5. Overview of Re-Glyco algorithm ... 16
4.6. GlcNAc scanning and AlphaFold glycoproteins test ... 18

**References** ... **19**

### List of Supplementary Figures

**Figure 1: A) Cumulative and individual explained variance by principal components. B) Silhouette score analysis for determining the optimal number of clusters. The plot depicts the silhouette scores for different numbers of clusters, with the peak at 6 clusters suggesting it as the optimal count for the given dataset. C) Comparison of centroids determined by Gaussian Mixture Model (GMM) and Kernel Density Estimation (KDE) in an asymmetric dataset. The scatter plot visualizes the data points with the GMM centroids (marked as 'X') and KDE centroids (marked as circles), showcasing the difference in centroid location due to the asymmetric distribution of the data. .............................. 8**

**Figure 2: Torsion Angles($\varphi, \psi$) between the protein($P$) and the glycan($G$). 12**

**Figure 3: Identified torsion pairs in a glycan($G$). .................................. 15**

**Figure 4: Re-Glyco algorithm flowchart ................................................ 17**

### List of Algorithms

### 1. GlycoShape Website

#### 1.1. Cross referenced data.

In order to complement the structural data represented within our resource, GlycoShape makes uses of other published resources to provide users with other supportive information. In particular, this additional information includes providing alternative naming formats for each glycan, chemical information for each glycan, and biological information for each glycan. This additional information is obtained from previously published glycoinformatics resources, namely GlyCosmos[1], GlyGen[2], and Glycowork[3]. The specific functions each of these resources is utilized for within the GlycoShape platform is discussed in Table 1. Furthermore, the use of many naming systems and GlyTouCan IDs lends the GlycoShape GDB to be compatible with previously established resources, and for it to be accessible from other resources itself.

TABLE 1. Overview of the glycoinformatic resources utilized within the GlycoShape platform. For each resource, a description of the information provided for the GlycoShape GDB is provided, along with the API address or python module that performs this function.

| GlycoShape GDB information | API service / Python module |
|---|---|
| Glycam to IUPAC name conversion | GlycoShape GAP |
| IUPAC to monosaccharide composition | GlycoShape GAP |
| IUPAC to WURCS name conversion | https://api.glycosmos.org/glycanformatconverter/2.8.2/iupaccondensed2wurcs |
| IUPAC to GlyTouCan ID fetching | https://api.glycosmos.org/glycanformatconverter/2.8.2/iupaccondensed2wurcs |
| GlyTouCan ID to biological information | https://api.glygen.org/glycan/detail/ |
| IUPAC to SNFG image creation | glycowork.motif.draw |
| IUPAC to motif identification | glycowork.motif.annotate |
| IUPAC to chemical information | glycowork.motif.annotate |
| IUPAC to termini identification | glycowork.motif.annotate |
| IUPAC to SMILES name conversion | glycowork.motif.processing |
| Canonicalize IUPAC name | glycowork.motif.processing |
| IUPAC to biological information | SugarBase from GlycoWork |

##### 1.2. **Frontend and APIs.**

In the development of the GlycoShape web application, a modern and efficient technology stack was employed to ensure robust performance and user-friendly interfaces. The frontend of the application is crafted using React TypeScript, a powerful JavaScript library for building user interfaces with TypeScript, which adds static type definitions to enhance code quality and understandability. For styling and theming, Chakra-UI, a simple, modular, and accessible component library, is utilized to create a consistent and responsive design across the application. The frontend build process is managed using npm@9.8.1 version, ensuring a streamlined and efficient compilation of the codebase.

The search feature of GlycoShape website uses Levenshtein Distance to calculate the differences & similarities between sequences and return the tops results. Also, the search by drawing glycan feature uses the SugarDrawer[4] tool which significantly enhances the user's ability to search for glycans, providing a more flexible and intuitive search experience.

On the backend, GlycoShape leverages a set of APIs developed using Flask, a lightweight and powerful web framework for Python. These APIs are responsible for fetching and managing the data displayed on the website. To ensure reliable and continuous service, the APIs are served using Gunicorn, a Python WSGI HTTP Server, behind an Nginx proxy server. This configuration offers enhanced performance, security, and scalability, making GlycoShape a reliable tool for concurrent users.

The latest details and documentation of the stable APIs are available at GlycoShape API Documentation `https://glycoshape.io/api-docs` . This documentation provides users and developers with comprehensive information on the API endpoints, usage, and response formats, facilitating easy integration and utilization of GlycoShape's capabilities in various other glyco-bioinformatics software and resources.

### 2. Free Glycan Simulation Protocol

The free glycan simulation systems were constructed using the Carbohydrate Builder tool of GLYCAM Web[5], with all glycans having a hydroxyl group at the terminal of the reducing sugar. This tool is capable of constructing multiple starting conformations of the glycan depending on the glycosidic angles being in the gauche-gauche, gauche-trans, or trans-trans conformation. For glycans that were predicted to have a one likely conformer from Carbohydrate Builder, triplicates simulations of 500 ns each were performed. For glycans that were predicted to have two likely conformers from Carbohydrate Builder, duplicates simulations of 500 ns were performed for each of the conformers. For glycans that were predicted to have three or more likely conformers from Carbohydrate Builder, a single replicate simulation of 500 ns was performed for each conformer. This guarantees that every glycan within GlycoShape has been simulated for a minimum of 1.5 µs. This long timescale of simulation ensures that the structural landscape of the glycan has been sufficiently sampled.

Each simulation system was constructed using the tleap module of AmberTools[6]. The systems were solvated in a water box with a minimum distance of between 12 Å and 15 Å, and ions were added to neutralize any system charges to a total concentration of between

150 mM and 200 mM NaCl. These values vary slightly between simulations, with the specific values available for each system on the Simulation Information tab for each glycan on the GlycoShape website. The GLYCAM06j-1[7] force field was used to model glycans, and the TIP3P water model was used to model solvent molecules[8]. For glycans with modifications not present in the GLYCAM06j-1 force field, such as the addition of phosphorylcholine groups as found in *Nematoda*, the General AMBER Force Field 2 (GAFF2)[9] force field was used to create additional parameters.

Simulations were then conducted using either AMBER 18[10] on resources provided by the Irish Centre for High-End Computing (ICHEC), or GROMACS 2022.4[11, 12]

For simulations conducted using AMBER, the energy of the system was minimized using the steepest descent algorithm, with all heavy atoms of the glycan restrained with a potential weight of 5 $kcal.mol^{-1}$. $Å^{-2}$. The system was then equilibrated in the NVT ensemble, with the system gradually heated from 0 to 100 K, and then from 100 K to 300 K. The system was then equilibrated in the NPT ensemble to maintain the pressure at 1 bar. The temperature was maintained at 300 K using Langevin dynamics with a collision frequency of 1 $ps^{-1}$, and the pressure was maintained at 1 bar using isotropic position scaling with a Berendsen barostat and a pressure relaxation time of 2 ps. Periodic boundary conditions were used throughout the simulations. The Van der Waals interactions were truncated at 11 Å and PME was used to treat long range electrostatics with B-spline interpolation of order 4. The SHAKE algorithm was used to constrain all bonds containing hydrogen atoms and to allow the use of a 2 fs time step for all simulations.

For simulations conducted using GROMACS, the prm7 and rst7 files produced by tleap were converted to GROMACS-readable top and gro file using ACPYPE[13]. The energy of the system was minimized using the steepest descent algorithm. The system was then equilibrated in the NVT ensemble at 300 K, and then equilibrated in the NPT ensemble to maintain the pressure at 1 bar. The temperature was maintained at 300 K using Langevin dynamics with a collision frequency of 1 $ps^{-1}$, and the pressure was maintained at 1 bar using anisotropic position scaling with a Parrinello-Rahman barostat and a pressure relaxation time of 5 ps. Periodic boundary conditions were used throughout the simulations. The Van der Waals interactions were truncated at 11 Å and PME was used to treat long range electrostatics. The LINCS algorithm was used to constrain all bonds containing hydrogen atoms and to allow the use of a 2 fs time step for all simulations.

### 3. Glycan Analysis Pipeline

The below section contains some key details in the GAP algorithm. We starts with merging individual MD trajectories from the uncorrelated replicas into one dataset and hydrogens are removed in the process to improve the computational effiencly.

#### 3.1. Molecule representation.

We first computes the flattened Euclidean distance matrix for a given set of data points. Given a dataset $X = \{x_1, x_2, ..., x_n\}$, where each $x_i$ in $\boldsymbol{R}^d$, the function computes the Euclidean distance matrix $G \in \boldsymbol{R}^{n\times n}$, where each element $g_{\mathrm{ij}}$ is given by:

$$g_{ij} = \left|\left|x_i - x_j\right|\right|_2 = \sqrt{\sum_{k=1}^{d} \left(x_{ik} - x_{jk}\right)^2} \tag{1}$$

The function then flattens the lower triangle of the distance matrix, excluding the diagonal. The flattened array $G_f$ is defined as follows:

$$G_f = \left\{g_{11}, g_{21}, g_{31}, ..., g_{n1}, g_{32}, ..., g_{\frac{n(n-1)}{2}}\right)\} \tag{2}$$

where each element in $G_f$ is obtained from the lower triangle of the distance matrix $G$. Finally, the function returns the transformed flattened array $G \in \boldsymbol{R}^{m\times p}$, where $p$ is the length of the flattened distance matrix $G_i$, and each row $i$ of $G$ contains the corresponding $G_i$.

#### 3.2. Dimentionality reduction.

Principal Component Analysis (PCA) is then applied to the matrix $G$ to reduce the dimensionality to a specified number of components $dim$ (= 3 for our case). The PCA transformation can be represented as follows:

$$T = G \times W \tag{3}$$

where $T \in \boldsymbol{R}^{m\times \mathrm{dim}}$ is the transformed data matrix, and $W \in \boldsymbol{R}^{p\times \mathrm{dim}}$ is the matrix containing the top `dim` principal components as columns.

The function returns the transformed data matrix $T$ in the form of a DataFrame. Additionally, the explained variance ratio and cumulative sum of the eigenvalues are calculated, which can be used to evaluate the explained variance of the PCA components as shown in figure Figure 1 A.

*dim* is set to three-dimension because as the number of dimensions increases, clustering efficiency tends to decrease. This is due to the fact that points become more distant from each other in higher-dimensional spaces.

##### 3.3. **Clustering algorithm and number of clusters.**

For clustering the conformational landscape Gaussian Mixture Model (GMM) is used as it presents distinct advantages over other clustering methods. GMM is a probabilistic approach that posits data generation from multiple Gaussian distributions

Module `mixture.GaussianMixture` from sklearn[14] used to carry out the Clustering with `covariance_type='full'`, and `random_state=42` for reproducibility.

The determination of the optimal number of clusters within GMM is guided by the silhouette score as shown in figure Figure 1 B. This metric evaluates how similar an object is to its own cluster compared to other clusters, thus ensuring the most coherent and distinct grouping of the conformational states.

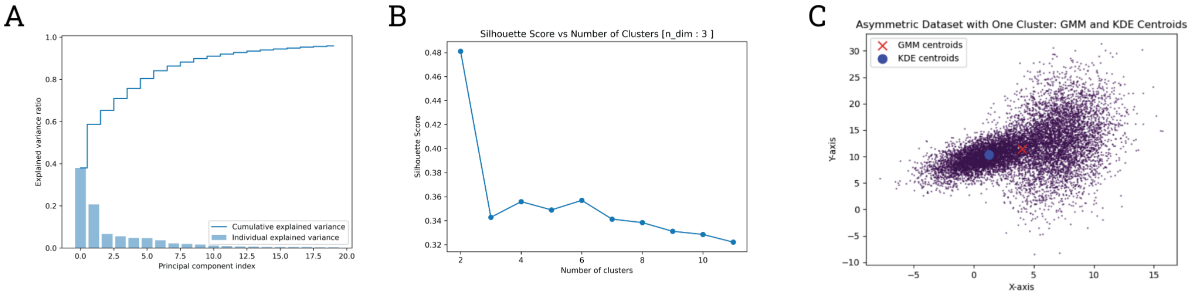

Figure 1. **A)** Cumulative and individual explained variance by principal components. **B)** Silhouette score analysis for determining the optimal number of clusters. The plot depicts the silhouette scores for different numbers of clusters, with the peak at 6 clusters suggesting it as the optimal count for the given dataset. **C)** Comparison of centroids determined by Gaussian Mixture Model (GMM) and Kernel Density Estimation (KDE) in an asymmetric dataset. The scatter plot visualizes the data points with the GMM centroids (marked as 'X') and KDE centroids (marked as circles), showcasing the difference in centroid location due to the asymmetric distribution of the data.

##### 3.4. **Representative structure from clusters KDE maxima.**

In our analysis, we sought to pinpoint the maxima points of a Kernel Density Estimation (KDE) which would serve as a proxy for the cluster centroids in a multi-dimensional feature space. To achieve this, we developed a computational strategy that comprises several steps.

The KDE is defined as:

$$\hat{p}(x) = \frac{1}{n \cdot h} \sum_{i=1}^{n} K \frac{x - x_i}{h} \tag{4}$$

Here, $K(\cdot)$represents the Gaussian kernel function, expressed as:

$$K(u) = \frac{1}{\sqrt{2\pi}} e^{-\frac{1}{2}u^2} \tag{5}$$

and $h$ denotes the bandwidth, a parameter that influences the smoothness of the density estimation. To determine the optimal bandwidth, we employ cross-validation, specifically a 5-fold cross-validation scheme, where the dataset is partitioned into five subsets. The model is trained on four subsets and validated on the fifth, iteratively. The bandwidth that minimizes the validation error across all folds is selected.

The search for the point nearest to the KDE's peak within each cluster begins by defining a negative KDE score function:

$$s(x) = -\hat{p}(x) \tag{6}$$

Minimizing ( s(x) ) equates to maximizing the KDE, thus pinpointing the density peak. For optimization, we utilize the L-BFGS-B algorithm, which allows for bound constraints —these constraints are based on the minimum and maximum values across each dimension of the cluster's data points.

Concretely, the process involves fitting a Gaussian Mixture Model to the data, assigning data points to the most probable cluster, and then, for the data points within a chosen cluster, computing the optimal bandwidth for KDE. With the KDE model established, we then apply the L-BFGS-B optimization method to identify the point that maximizes the KDE, which corresponds to the cluster's most representative or central point. This approach is particularly useful for elucidating the conformational landscape of molecular clusters, where traditional centroid-finding methods may not be applicable due to the irregularity and asymmetry of data distributions as shown in Figure 1 C.

### 4. Re-Glyco: a glycoprotein builder

This section contains key ideas involed in development of Re-Glyco tool. All the parameters is selected based on compute time and results accuracey.

#### 4.1. **Glycoprotein torsion information.**

To re-glycosylate proteins with glycan structures from GlycoShape, it is necessary to know the dihedral angles between the reducing sugar and the sidechain of the protein sidechain. For N-glycosylation, the dihedral angles of the glycosidic linkages between the terminal GlcNAc residue and the Asn sidechain were obtained from the Privateer package10. However, for O-glycosylation it is necessary to obtain these angles from MD simulations of exemplar proteins for each glycosylation type, as shown in Table 2

The simulation systems of syndecan-1[15] and thrombospondin repeat 2 were made using the tleap module of AmberTools[6]. The simulation systems of 3-phosphoinositide-dependent protein kinase 1[16], epidermal-like growth factor from coagulation factor IX[17, 18], and dystroglycan 1[19] were made using the Solution Builder module of CHARMM-GUI[20]. The charged N- and C-terminal residues were neutralized by capping with ACE and NME groups, respectively. The systems were solvated in a water box with a minimum distance of 12 Å, and ions were added to neutralize any system charges to a total concentration of 150 mM NaCl. The AMBER 14SB force field[21] was used to model proteins and ions, the GLYCAM06j-1[7] force field was used to model glycans, and the TIP3P water model was used to model solvent molecules4.

The glycophorin A transmembrane system was made using the Membrane Builder module of CHARMM-GUI[22]. The charged N- and C-terminal residues were neutralized by capping with ACE and NME groups, respectively. The protein was embedded in a 130 x 130 Å POPC membrane using the PPM v2.0 server[23], and ions were added to neutralize any system charges to a total concentration of 150 mM NaCl. The AMBER 14SB force field[21] was used to model proteins and ions, the GLYCAM06j-1[7] force field was used to model glycans, the Slipids[24] force field was used to model lipid molecules, and the TIP3P water model was used to model solvent molecules[8].

Table 2. Summary of MD simulations conducted to obtain dihedral angles between the glycosidic bonds of the reducing sugar and the glycosylated protein sidechain. For each simulation system, the PDB ID is given where an experimentally-determined protein structure was simulated, and a UniProt ID where an AlphaFold structure was simulated.

| **Protein** | **PDB/ UniProt ID** | **Glycosylated residue(s)** | **Glycosylation types (s)** | **Glycan(s)** | **Simulated time (μs)** |
|---|---|---|---|---|---|
| Thromospondon repeat 2 | O14514 | T369 | O-Fucose | Glc(b1-3)Fuc(a1- | 1 |
| Glycophorin A | P02724 | S63, S66, and T69 | O-GalNAc | Neu5Ac(a2-3)-Gal(b1-3) [Neu5Ac(a2-6)]GalNAc(a1- | 0.75 |
| 3-phosphoinositide-dependent protein kinase 1 | 1H1W | S92, S105, T222 | O-GlcNAc | GlcNAc(b1- | 1 |
| Epidermal-like growth factor from Coagulation Factor IX | 1EDM | S99 and S107 | O-Glucose and O-Fucose | Glc(b1- and Fuc(a1- | 1 |
| Epidermal-like growth factor from Coagulation Factor IX | 5VYG13 | S99 | O-Glucose | Xyl(a1-3)Xyl(a1-3) Glc(b1- | 1 |
| Dystroglycan 1 | 5GGP | T319 | O-Mannose | GlcNAc(b1-2)-Man(a1- | 1 |
| Syndecan-1 | 6EJE | S206 | O-Xylose | GlcA(b1-3)GalNAc4S(b1-4) GlcA(b1-3)GalNAc4S(b1-4) GlcA(b1-3)-Gal(b1-3)-Gal(b1-4)Xyl(b1-) | 1 |

For all soluble protein systems, the energy of the system was minimized using the steepest descent algorithm, with all heavy atoms of the protein and glycan restrained with a potential weight of 5 kcal.mol$^{-1}$ .Å$^{-2}$. The system was then equilibrated in the NVT ensemble, with the system gradually heated from 0 to 100 K, and then from 100 K to 300 K. The system was then equilibrated in the NPT ensemble to maintain the pressure at 1 bar. The temperature was maintained at 300 K using Langevin dynamics with a collision frequency of 1 ps-[1], and the pressure was maintained at 1 bar using isotropic position scaling with a Berendsen barostat and a pressure relaxation time of 2 ps. Periodic boundary conditions were used throughout the simulations. The Van der Waals interactions were truncated at 11 Å and Particle Mesh Ewald (PME) was used to treat long range electrostatics with B-spline interpolation of order 4. The SHAKE algorithm was used to constrain all bonds containing hydrogen atoms and to allow the use of a 2 fs time step for all simulations.

For the transmembrane system, the energy of the system was minimized using the steepest descent algorithm, with all atoms of the protein, lipid head groups, and glycans restrained. The system was then equilibrated in the NVT ensemble, with the system gradually heated from 0 to 100 K, and then from 100 K to 300 K. The system was then equilibrated in the NPT ensemble to maintain the pressure at 1 bar. Position restraints placed on the atoms of the protein, lipid head groups, and glycans were gradually released during the equilibration process. The temperature was maintained at 300 K using Langevin dynamics with a collision frequency of 1 ps-[1], and the pressure was maintained at 1 bar using semi-isotropic position scaling with a Berendsen barostat and a pressure relaxation time of 1 ps. Periodic boundary conditions were used throughout the simulations. The Van der Waals interactions were truncated at 11 Å and Particle Mesh Ewald (PME) was used to treat long range electrostatics with B-spline interpolation of order 4. The SHAKE algorithm was used to constrain all bonds containing hydrogen atoms and to allow the use of a 2 fs time step for all simulations.

All simulations were performed using AMBER18[10] on resources provided by the Irish Centre for High-End Computing (ICHEC).

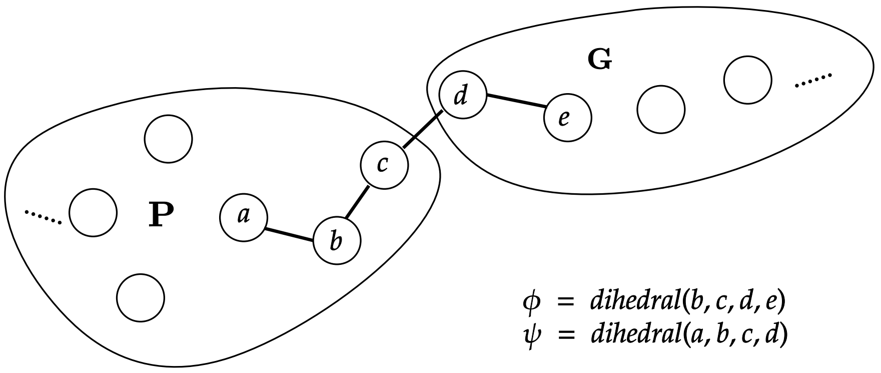

FIGURE 2. Torsion Angles($\varphi, \psi$) between the protein($P$) and the glycan($G$).

Currently supported function includes N-GlcNAcylation, O-GalNAcylation, O-GlcNAcylation, O-Fucosylation, O-Mannosylation, O-Glucosylation, O-Xylosylation, C-Mannosylation

TABLE 3. Torsion ranges for different glycosylations

| **Residue** | **Sugar** | **Φ Angles** | **Ψ Angles** | **Link Type** |
|---|---|---|---|---|
| ASN | GlcNAc | (-130, -63) [CG, ND2, C1, O5] | (152, 205) [CB, CG, ND2, C1] | beta |
| THR | GalNAc | (55, 83) [CB,OG, C1, O5] | (86, 142) [CA, CB, OG, C1] | alpha |
| THR | Fuc | (274, 304) [CB,OG, C1, O5] | (148, 182) [CA, CB, OG, C1] | alpha |
| THR | Man | (61, 88) [CB,OG, C1, O5] | (88, 150) [CA, CB, OG, C1] | alpha |
| THR | GlcNAc | (143, 221) [CB,OG, C1, O5] | (171, 193) [CA, CB, OG, C1] | alpha |
| SER | GalNAc | (61, 86) [CB, OG, C1, O5] | (163, 224) [CA, CB, OG, C1] | alpha |
| SER | Fuc | (58, 106) [CB, OG, C1, O5] | (86, 192) [CA, CB, OG, C1] | alpha |
| SER | Glc | (261, 297) [CB, OG, C1, O5] | (146, 219) [CA, CB, OG, C1] | beta |
| SER | Xyl | (265, 309) [CB, OG, C1, O5] | (107, 231) [CA, CB, OG, C1] | beta |
| SER | GlcNAc | (272, 308) [CB, OG, C1, O5] | (178, 300) [CA, CB, OG, C1] | beta |
| TRP | Man | (110, 150) [CG, CD1, C1, O5] | (-3, 3) [CB, CG, CD1, C1] | alpha |

##### 4.2. $P$, $G$ and fitness function $F_{\{P,G,\varphi,\psi\}}$.

**ALGORITHM 1**. Steric Interaction Scoring Function

Steric Interaction Calculation

**Steric** $(Garr, Parr)$:

**initialize** $r$ **to** 0
**calculate** euclidean distances $D$ between $G$ **and** $P$ using `cdist`
select distances $C$ from $D$ where distance $< 1.7$
**for** each distance $i$ **in** C[2:−1]**:** // Ignoring H & "d"
    increment $r$ by $200 \times \exp(-i^2)$
**return** $r$

**Algorithm 2**. Fitness Function for Evaluating Steric Interactions Post-Rotation

Fitness Function

**Fitness F** ( $P$, $G$, $\varphi$,$\psi$):

**define** function `rotate` with parameters $G$, $\varphi$, $\psi$, $a$, $b$, $c$, $d$
**apply** `rotate` **to** $G$ **to** obtain $G'$
**invoke** `Steric` function with $G'$ **and** $P$ **to** compute steric interactions
**assign** the output of `Steric` function **to** variable **steric**
**return steric**

**Algorithm 3**. Algorithm for Rotating Molecular Group G Using Quaternion Rotation Matrices

Group Rotation Using Quaternion Matrices

**rotate** ($G$, $phi$, $psi$, $a$, $b$, $c$, $d$):

**initialize** $q_{\{\varphi\}}$ as quaternion representing rotation by $\varphi$ around axis $b-c$
**initialize** $q_{\{\psi\}}$ as quaternion representing rotation by $\psi$ around axis $c-d$
**for** each atom $x$ **in** $G$**:**
    **calculate** new position $x'$ by applying $q_{\{\varphi\}}$ **then** $q_{\{\psi\}}$ **to** $x$
    **update** position of $x$ **in** $G$ **to** $x'$
**return** modified $G$ as $G'$

##### 4.3. Wiggle algorithm for alternate conformation within a cluster.

The provided Figure 3 illustrates an algorithm designed to identify glycosidic bonds and the atoms involved within a molecular structure. This algorithm functions by pinpointing torsion pairs that facilitate the molecule's ability to undergo conformational changes. Initially, it identifies all the atoms forming part of cyclic structures within the molecule and isolates them for further examination. Subsequently, the algorithm scans through each atom in the non-cyclic parts of the molecule, singling out those with only two bonds (i.e., a degree of two). For these atoms, it identifies adjacent atoms and establishes torsion pairs, which are the quartets of atoms that define the axis of rotation for potential torsional movements. The collection of these torsion pairs effectively maps out all possible sites where the molecule can 'wiggle' or rotate, thus providing alternate conformation of the glycan within a cluster.

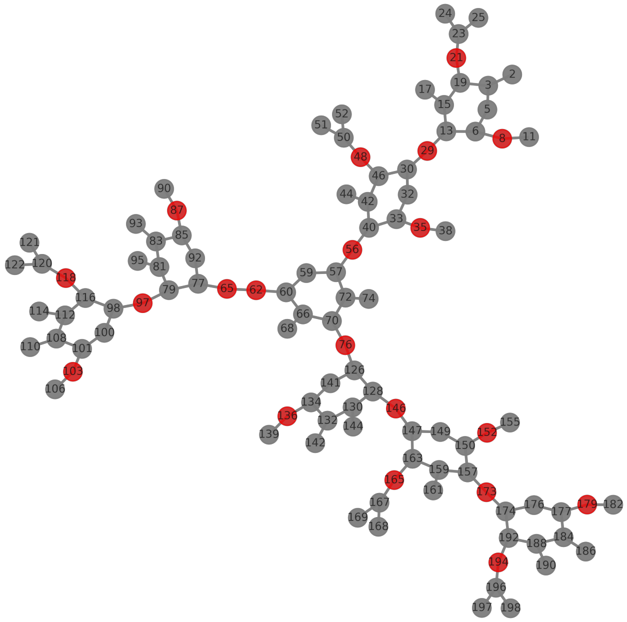

FIGURE 3. Identified torsion pairs in a glycan($G$).

**ALGORITHM 4**. Torsion Pairs Identification Algorithm

Torsion Pairs Identification in Molecular Structures

**torsion_pairs** ($G$, *connect*):

**let** empty list cycles
**for** each cycle **in** nx.cycle_basis(G)**:**
    **return** add atoms **in** cycle **to** cycles
**let** empty list pairs
**for** each node **in** “G”**:**
    **if** degree of node $==$ 2 **and** node **not in** cycle**:**
        find neighbors $j, k$ of node
        **if** degree of $j > 1$**:**
            find neighbors l of j excluding node
            pairs.append([$l$[0], $j$, node, $k$])
        **if** degree of $k > 1$**:**
            find neighbors $m$ of $k$ excluding node
            pairs.append([$m$[0], $k$, node, $j$])
**return** pairs

**Algorithm 5**. Algorithm for Generating New Conformations by Random Torsion Rotation

Random Torsion Rotation for Molecular Conformation

**wiggle** ($G$, *pairs*):

**let** angle_range be (-10, 10) degrees
**let** $G_{\text{new}}$ be a copy of G
**for** each pair **in** pairs: // from torsion_pairs
    **let** [$a$, $b$, $c$, $d$] be atoms **in** the torsion pair
    **let** random_angle be a random value from angle_range
    rotate $G_{\text{new}}$ at torsion defined by [$a$, $b$, $c$, $d$] by random_angle using quaternion rotation
**return** $G_{\text{new}}$

##### 4.4. Genetic optimization algorithm for finding optimal $\varphi$ and $\psi$.

**Algorithm 6**. Genetic Algorithm for Optimization of Molecular Conformations

Genetic Algorithm for Minimizing Fitness Function

**genetic_algorithm** ($P$, $G$, $[\varphi_{\text{range}}]$, $[\psi_{\text{range}}]$):

**let** population_size be 128
**let** mutation_rate be 0.2
**let** generations be 8
**let** individial$_{\text{best}}$ be $\varphi_{\text{best}}$ **and** $\psi_{\text{best}}$ with fitness$_{\text{best}}$ 100000
**let** population be an empty list of size population_size
**for** each individual **in** population:
    **let** $\varphi_{\text{individual}}$ be a random number within $[\varphi_{\text{range}}]$
    **let** $\psi_{\text{individual}}$ be a random number within $[\psi_{\text{range}}]$
**for** gen from 1 **to** generations:
    **for** each individual **in** population:
        **let** individual [fitness] be F(P, G, $\varphi_{\text{individual}}$, $\psi_{\text{individual}}$)
    **sort** population by increasing fitness
    **let** parents be the top 50% of population
    **let** children be crossover_and_mutate(parents, mutation_rate)
    replace_worst_with_children(population, children)
    **if** fitness of the first individual **in** the sorted population < fitness$_{\text{best}}$
        **let** best$_{\text{individial}}$ be the first individual **in** the sorted population
**return** individial$_{\text{best}}$ ($\varphi_{\text{best}}$ **and** $\psi_{\text{best}}$)

##### 4.5. Overview of Re-Glyco algorithm.

Re-Glyco begins with the major cluster (cluster0) and seeks $\varphi$ and $\psi$ angles within permissible ranges derived from MD simulations or crystal structures. The goal is to minimize steric hindrance, defined by a specific loss function $F(P, G, \varphi, \psi)$, over a population of 128 and across 8 generations for each glycan conformation. If steric clashes persist, Re-Glyco shifts to the second-best major conformation from the GlycoShape database and so on. If no conformation fits the protein site, Re-Glyco tests the least steric glycan conformation, repeating the process (up to 40 times) with adjustments ("wiggles") involving random $10^o$ rotamer changes.

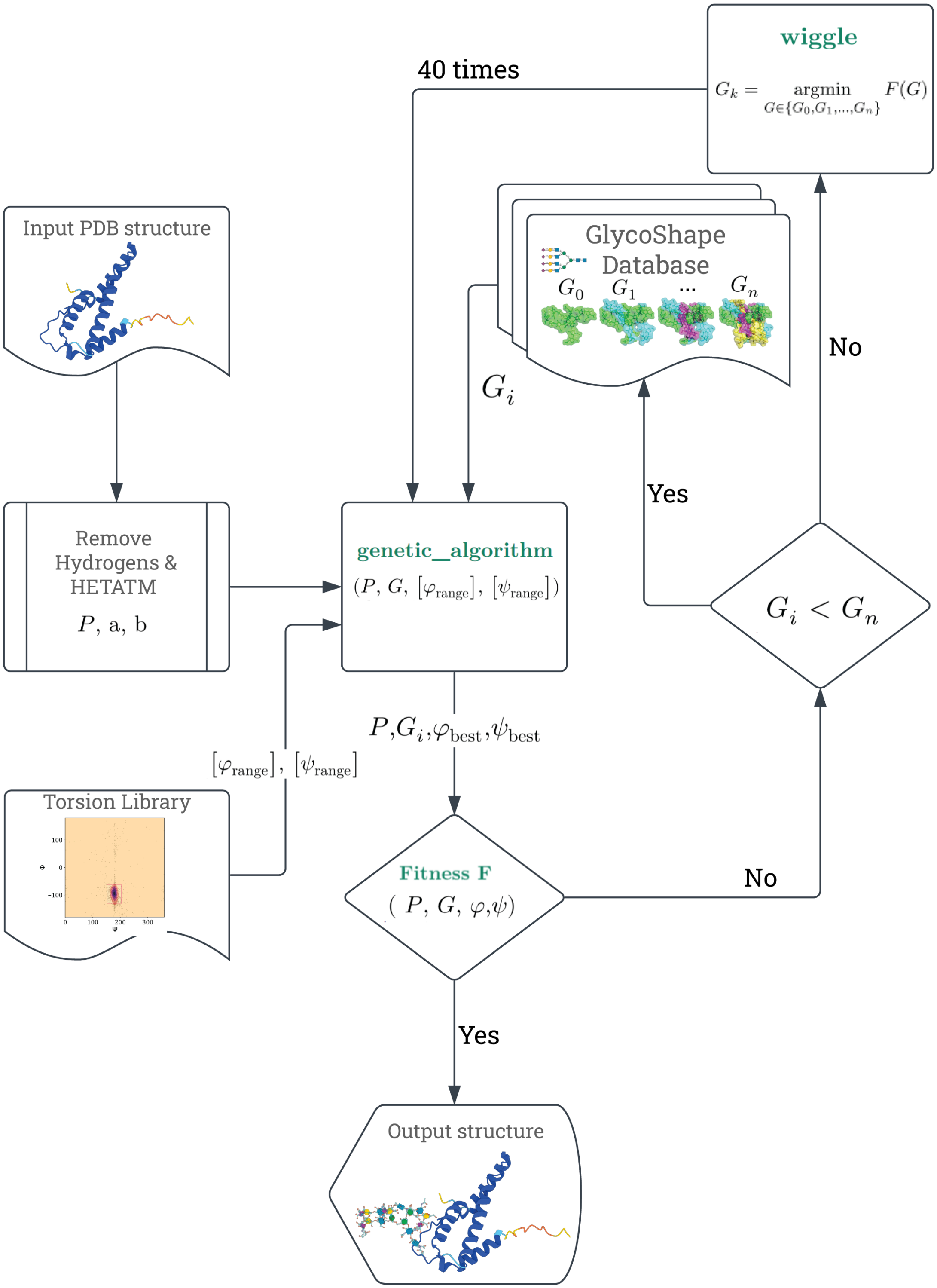

FIGURE 4. Re-Glyco algorithm flowchart

#### 4.6. GlcNAc scanning and AlphaFold glycoproteins test.

The GlcNAc Scanning and AlphaFold test data have been made available in JSON format at the GlycoShape GitHub repository. The test datasets for GlcNAc Scanning include detailed structural information such as the UniProt ID, amino acid sequence, and specific glycosylation sites with their solved clashes, torsion angles, and cluster assignments. Similarly, the AlphaFold test data provide insights into the computational time, clash resolution status, predicted glycosylation sites with their respective dihedral angles, and the per-residue prediction confidence scores (pLDDT) from AlphaFold's models.

`https://github.com/Ojas-Singh/GlycoShape/tree/main/GlcNAc_Scanning_and_AF_test_data`

Table 4. Default glycan list for uniport annotated PTM information

| Glycosylation Type | Glycan Composition |
|---|---|
| C-linked (Man) tryptophan | Man |
| O-linked (Fuc...) serine | Fuc |
| O-linked (Fuc...) threonine | Fuc |
| O-linked (GalNAc...) serine | Neu5Ac(a2-3)Gal(b1-3)GalNAc |
| O-linked (GalNAc...) threonine | Neu5Ac(a2-3)Gal(b1-3)GalNAc |
| O-linked (Glc...) serine | Glc |
| O-linked (Glc...) threonine | Glc |
| O-linked (Man...) serine | Neu5Ac(a2-3)Gal(b1-4)[Fuc(a1-3)]GlcNAc(b1-2)Man |
| O-linked (Man...) threonine | Neu5Ac(a2-3)Gal(b1-4)[Fuc(a1-3)]GlcNAc(b1-2)Man |
| O-linked (Xyl...) serine | Xyl |
| O-linked (Xyl...) threonine | Xyl |
| O-linked (Xyl...) (chondroitin sulfate) serine | GalNAc(b1-4)GlcA(b1-3)GalNAc(b1-4)GlcA(b1-3)-GalNAc(b1-4)GlcA(b1-3)GalNAc(b1-4)GlcA(b1-3)-Gal(b1-3)Gal(b1-4)Xyl |
| O-linked (Xyl...) (chondroitin sulfate) threonine | GalNAc(b1-4)GlcA(b1-3)GalNAc(b1-4)GlcA(b1-3)-GalNAc(b1-4)GlcA(b1-3)GalNAc(b1-4)GlcA(b1-3)-Gal(b1-3)Gal(b1-4)Xyl |
| O-linked (GlcNAc) serine | GlcNAc |
| O-linked (GlcNAc) threonine | GlcNAc |
| N-linked (GlcNAc...) asparagine | GlcNAc(b1-2)Man(a1-3)[GlcNAc(b1-2)Man(a1-6)]-Man(b1-4)GlcNAc(b1-4)GlcNAc |
| N-linked (GlcNAc...) (complex) asparagine | Neu5Ac(a2-6)Gal(b1-4)GlcNAc(b1-2)Man(a1-3) [Neu5Ac(a2-6)Gal(b1-4)GlcNAc(b1-2)Man(a1-6)]-Man(b1-4)GlcNAc(b1-4)[Fuc(a1-6)]GlcNAc |
| N-linked (GlcNAc...) (hybrid) asparagine | Neu5Ac(a2-6)Gal(b1-4)GlcNAc(b1-2)Man(a1-3) [Man(a1-3)[Man(a1-6)]Man(a1-6)]Man(b1-4)GlcNAc(b1-4)GlcNAc |
| N-linked (GlcNAc...) (high mannose) asparagine | Man(a1-3)[Man(a1-6)]Man(a1-6)[Man(a1-3)]-Man(b1-4)GlcNAc(b1-4)GlcNAc |
